# A low-cost and open-source solution to automate imaging and analysis of cyst nematode infection assays for *Arabidopsis thaliana*

**DOI:** 10.1101/2022.07.14.500020

**Authors:** Olaf Prosper Kranse, Itsuhiro Ko, Roberta Healey, Unnati Sonawala, Siyuan Wei, Beatrice Senatori, Francesco De Batté, Ji Zhou, Sebastian Eves-van den Akker

## Abstract

**Background:** Cyst nematodes are one of the major groups of plant-parasitic nematode, responsible for considerable crop losses worldwide. Improving genetic resources, and therefore resistant cultivars, is an ongoing focus of many pest management strategies. One of the major bottlenecks in identifying the plant genes that impact the infection, and thus the yield, is phenotyping. The current available screening method is slow, has unidimensional quantification of infection limiting the range of scorable parameters, and does not account for phenotypic variation of the host. The ever-evolving field of computer vision may be the solution for both the above-mentioned issues. To utilise these tools, a specialised imaging platform is required to take consistent images of nematode infection in quick succession.

**Results:** Here, we describe an open-source, easy to adopt, imaging hardware and trait analysis software method based on a pre-existing nematode infection screening method in axenic culture. A cost-effective, easy-to-build and -use, 3D-printed imaging device was developed to acquire images of the root system of *Arabidopsis thaliana* infected with the cyst nematode *Heterodera schachtii*, replacing costly microscopy equipment. Coupling the output of this device to simple analysis scripts allowed the measurement of some key traits such as nematode number and size from collected images, in a semi-automated manner. Additionally, we used this combined solution to quantify an additional trait, root area before infection, and showed both the confounding relationship of this trait on nematode infection and a method to account for it.

**Conclusion:** Taken together, this manuscript provides a low-cost and open-source method for nematode phenotyping that includes the biologically relevant nematode size as a scorable parameter, and a method to account for phenotypic variation of the host. Together these tools highlight great potential in aiding our understanding of nematode parasitism.

## Introduction

Plant-parasitic nematodes represent a fraction of the total number of free-living nematode species [1], but are widely studied due to their agricultural importance: their parasitism accounts for over 10 % of the annual life-sustaining crop losses, costing the industry roughly 100-157 billion U.S. dollars per year [1–3]. In some cropping systems, plant-parasitic nematodes are the dominant plant pathogen of any kind [4].

Cyst nematodes are one of the major groups of plant-parasitic nematode, infecting a variety of crops globally. In general, the cyst nematode life cycle is as follows; a cyst, dormant in the soil for potentially decades, containing hundreds of eggs, is activated by the presence of diffusates from roots growing nearby. From these eggs, the infective second-stage juvenile (J2) hatches. The J2 migrates towards the growing root, penetrates host cells using a rapid thrusting action of their needle-like stylet, through which they secrete a variety of proteins to facilitate parasitism [5]. In the vasculature of the plant, cyst nematodes stimulate the formation of a “feeding-site” by re-differentiating existing procambial or pericyclic cells [6–8]. Feeding sites are unique pseudo-organs, unlike any other plant tissue, and are characterised by reduced and fragmented vacuoles, an increased smooth endoplasmic reticulum, and proliferated ribosomes, mitochondria, and plastids [9–11]. These organs are formed by partial cell wall dissolution and subsequent protoplast fusion of hundreds of adjacent cells, giving rise to a large, multi-nucleate, syncytial cell from which the nematode derives all external nutrition.

These obligate biotrophs will remain attached to this single feeding site for several weeks, maintaining a prolonged interaction with living host tissue. As the nematode feeds its body swells and increases in size dramatically. The sex of the nematode is determined epigenetically, during parasitism, and is linked to food availability. If the feeding site does not provide sufficient nutrition, the nematode terminally develops into a male, stops feeding, regains mobility, hatches out of the J3/4 body, and leaves the plant tissue for mating. Fertilised females produce eggs within their body, the body wall tans and hardens, and she falls off the plant leaving a cyst containing eggs that can live for years within the soil [12].

Current control measurements focus on the reduction of the density of plant-parasitic nematodes in the field and minimising the spread to other agricultural lands. Keeping the nematode population under a “damage threshold” is mainly achieved by crop rotation and the use of resistant cultivars and/or pesticides. However, in larger population densities, the rotation scheme may become economically unviable. Additionally, the use of the most effective nematicides is restricted due to concerns about the effects on human health and the environment. Improving genetic resources and developing impactful cultivars is therefore a current and future focus of many pest management strategies for plant-parasitic nematodes.

Plant genes that impact the outcome of nematode infection are varied. As with all plant-pathogens, classical resistance genes (NLR-type) have been identified against plant-parasitic nematodes, but in general our understanding of resistance remains limited due to the slow nature of screening for new ones. One of the major bottlenecks in identifying the plant genes that impact the outcome of infection is phenotyping. This is due in large part to the root-parasitic nature of these pathogens and the requirement of manual screening.

The most accessible infection assay is based on a well-established *H. schachtii:Arabidopsis thaliana* tissue culture system [13]. In brief, plants are grown in sterile tissue culture and infected about four weeks after germination. The total number of individuals infecting each plant is scored by manual assessment under the microscope. While extremely powerful, the assay has some clear limitations: i) it requires access to microscopy equipment; ii) due to the manual effort involved only a small number of genotypes can be screened at a time; and iii) it does not account for the phenotypic variation of the infectable tissue of the host.

There are limited examples in the literature reporting nematode phenotyping and associated phenotypic analysis using sensors such as red-green-blue (RGB) cameras [14–16]. However, some work on open-source hardware design for programmable imaging in plant research is reported. For example, PhenoBox [17], SeedGerm [18], CropSight [19], LiDARPheno [19], and HyperScanner [21]. In all these tools, a Raspberry Pi was used in combination with a light sensor to capture images of the subjects of interest. For analysing acquired images, open solutions such as Leaf-GP [22] using Python-based analytic packages, phenoSEED [23], AutoRoot [23], and PYM [25] using ImageJ/FIJI macro scripts to perform object segmentation and trait analysis. Taken together, the above-mentioned examples clearly illustrate a scope for development of a hardware/software solution for screening plant parasitic nematode infection.

In this paper we describe a low-cost easy to build and adopt image analysis approach to measure nematode infection. We describe methods to: 3D print and construct an “imaging tower”; capture images of infected plants in the well-established tissue culture system; use image analysis scripts to isolate, count, and measure parasitic nematodes; and normalise these data to the available root surface area at time of infection.

## Materials and methods

### Cultivation of nematodes and setting up the infection assay

To culture *Heterodera schachtii* on *Sinapis alba* (cv albatross) and for the infection assay on *Arabidopsis thaliana* (either Col-0 or N804585), seeds were surface sterilised with 20% dilution of 3.6 % sodium hypochlorite (ParoZone, Henkel) for 20 minutes and washed six times with sterile double distilled H_2_O. The seeds were kept at 4°C overnight to improve and synchronise germination [26]. The seeds were sown on sterile standard KNOP-medium (Duchefa Biochemie) [27] *in vitro* in a 15 cm sterile Petri dish (SARSTEDT) for *S. alba* and 5 cm deep well petri dishes (Thermo-Fisher) for *A. thaliana*. For purposes of female counting the agar was dyed using various food colouring (Limino, DYL-ghm-20210126-324) as described in Table 1. The different colours were tested in two different concentrations, a darker and a lighter variant. For purposes of cyst counting the agar remained undyed.

**Table 1:**
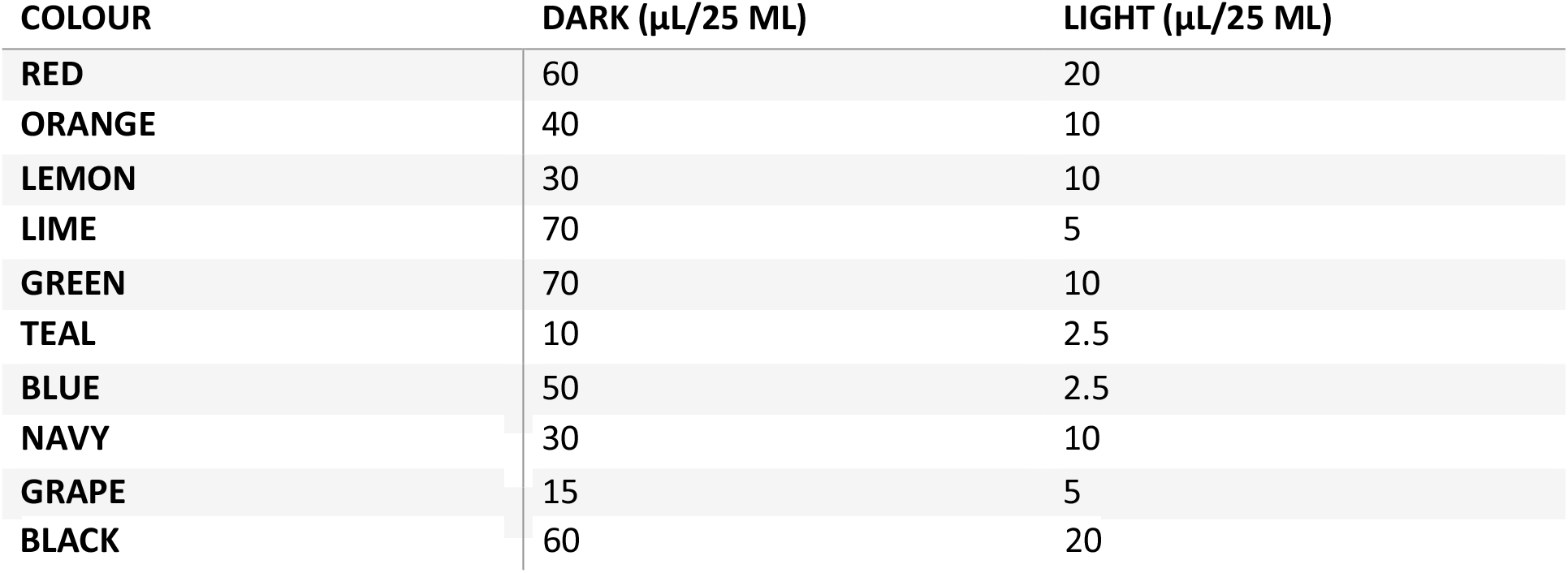
Amount of food colouring used in µL per 25 mL of KNOP-medium. In the first column the colour name as described by the manufacturer (Limino). The remaining two columns describe the different amounts of dye used per variant.

Plants grew on a 16 hour day 21°C and 8 hour night 20°C cycle in an MLR-352-PE growth chamber (Panasonic). Cysts were then soaked in 3 mM Zinc Chloride (SIGMA-ALDRICH) to promote hatching in a specialised hatching jar (Jane Maddern Cosmetic, 250mL) with two 2.5 cm plastic rings (alt-intech Tube Perspex) holding a 20 µm mesh (SIGMA-ALDRICH). Five days after hatching, J2 nematodes that had passed through the mesh were collected by pipetting. The density of nematodes was adjusted to 1 nematode/µL using sterile double distilled H_2_O containing 0.01% v/v Tween (Sigma-Aldrich). Once the plants reached an age of 21-28 days, the roots were inoculated with ∼300 J2 nematodes on *S. alba* and 26 days with ∼80 J2 nematodes on *A. thaliana* by pipetting the suspension on the roots. On *S. alba* it took 10-12 weeks at 20-25°C in darkness for the cysts to develop. Images of infected *A. thaliana* plates were taken depending on the need at least on two weeks post germination for quantification of root surface area, 21 days for female nematodes and 21+ days for encysted nematodes.

### Building hardware for image requirements

#### Open hardware design

Images of petri-dishes were taking using a custom imaging tower. The imaging components of the device are assembled onto a custom 3D printed apparatus (https://github.com/OlafKranse/A_low_cost_imaging_tower/tree/main/STL files. The STL files were sliced using CURA (Ultimaker) with an infill of 20%, a layer height of 0.15 mm, a wall thickness of 1.2 mm and a printing speed of 40.0 mm/s. The nozzle temperature was set to 205 °C and the build plate to 60 °C. The two components of the tower were printed in tough PLA with PVA as support (Ultimaker) in an Ultimaker S3 equipped with an 0.4 mm AA and BB core. The PVA was dissolved for 24 hours in tap water. The RPI-HQ-CAMERA (RASPBERRY-PI) was mounted in the holes of the top part of the tower using M2 bolts (2 cm), nuts, and washers (RS components) and the 16 mm Telephoto Lens (RASPBERRY-PI) was bolted onto the camera as instructed in Figure 1. The camera was connected to the Raspberry Pi via a 30 cm Ribbon Cable (THE PI HUT). Lastly, the 60 LED RGB STRIP LIGHT 6400K SET IP65 12V (V-TAC) was slid through the extrusions on the side of the bottom part (Figure 1).

**Figure 1:**
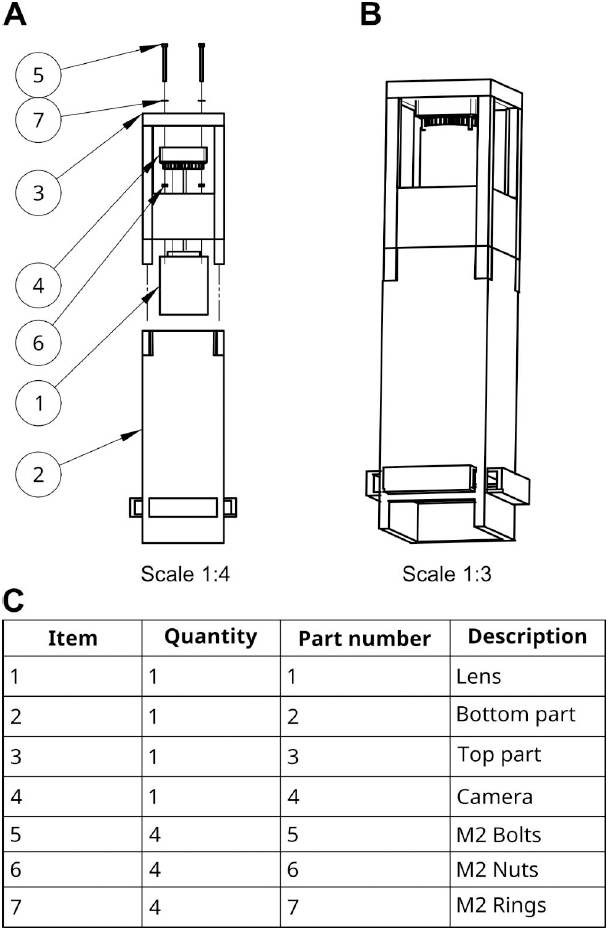
Explosion diagram of the imaging tower. A) The separate components labelled with their corresponding end position. The item numbers indicate a position in the table below. B) The final product after assembly. C) A table of components. The item number corresponds with the numbers in A.

#### Data collection and calibration

Before inoculation (four weeks post germination), images were taken using a custom script controlling the tower described above (https://github.com/OlafKranse/A_low_cost_imaging_tower/blob/main/ImagingandAnalyses/Imagingcommand.txt). Petri dishes were wiped clean using disposable wipes (Kimwipes) before capture. The images were separated into folders corresponding to the mutant line. Using ImageJ commands [28], the script, in order: converted the image to 8bit, subtracted the background, enhanced the contrast, ran a threshold, converted the output to a mask, performed a watershed, defined an area of interest, analysed and counted the resulting overlay using the parameters as described in the script (https://github.com/OlafKranse/A_low_cost_imaging_tower/blob/main/ImagingandAnalyses/automated_root_surface_area.ijm). A slightly adjusted script was used for plates containing dye (https://github.com/OlafKranse/A_low_cost_imaging_tower/blob/main/ImagingandAnalyses/automated_root_surface_area_colored_agar.ijm). The root surface area for all the images in the folder were exported as a CSV file. By adding colour thresholding to the above mentioned scripts, leaf surface area was isolated from images (lhttps://github.com/OlafKranse/A_low_cost_imaging_tower/blob/main/ImagingandAnalyses/Leaf_surface_count.ijm).

### Traditional counting and automatic counting

#### Ground truth by manual scoring

The number of females and males in manual counting was scored under a S9D Stereomicroscope (Leica). For manual size quantification an image was taken using the above-described imaging machine and an outline was drawn using the polygon selections tool in ImageJ (Figure 2). Using the measure command, the area within the outline was quantified in number of pixels.

**Figure 2:**
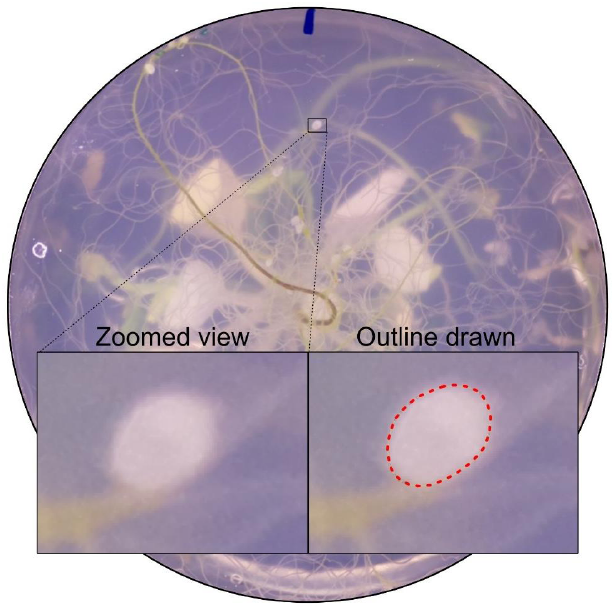
An example of a manual outline drawn around a female cyst nematode using ImageJ. Left) before drawing an outline. Right) the outline drawn.

#### Script-based counting and quantifiable traits

Automatic counting was performed on images taken as described above. Depending on the treatment a different script was used to calculate the number and size of females. Before isolation, the colour histogram for all images was normalised to the first image in the dataset using a custom python script (https://github.com/OlafKranse/A_low_cost_imaging_tower/tree/main/Imaging and Analyses/Normalise colour). The images were then processed in ImageJ for two different nematode life stages: i) tanned cyst nematodes; ii) female nematodes.

i. Automatic counting of cysts. Using ImageJ commands, the script, in brief: thresholded colour, converted the image to 8bit, removed pixel outliers, performed a watershed, waited for the user to define an area of interest, analysed and counted the resulting overlay using the parameters as described in the script (https://github.com/OlafKranse/A_low_cost_imaging_tower/tree/main/Imaging and Analyses).
ii. Semi-automatic counting of females. The well-established watershedding addon MorphoLibJ [29] was used for segmentation of female nematode from images of coloured agar. The tool parameters were set to a morphological gradient type and a watershed segmentation threshold both with a radius of 5. The display type was set to catchment basins and the number and size of segmentation was scored using the analyse regions function.

## Results

Cyst nematodes are naturally root parasitic obligate biotrophs, and notoriously difficult to phenotype during infection. Their ability to infect depends, although not exclusively, on a variety of host factors (including genotype and physiology), for example, root surface area. To account for this, custom hardware (low-cost, 3D printed imaging platform) and associated custom software (simple image analysis pipelines) were developed to standardise/semi-automate root and nematode phenotyping in the model *Heterodera schachtii*:*Arabidopsis thaliana* pathosystem.

### A cost-effective, open-source 3D printable imaging device

Based on *in vitro* 5 cm petri dish cultures of individual *A. thaliana* plants on transparent medium [13], a custom 3D printed imaging platform was designed to capture consistent images of the root systems. Consistency between images is essential for downstream image analysis (object segmentation using thresholding). The apparatus comprises of two halves (Figure 3A): the upper half housing a high-quality 12-megapixel camera and lens (Figure 3B): the bottom half housing an LED strip wrapping around the base and passes through the tower (Figure 3D). Combined with a Raspberry Pi computer, the setup can consistently capture high quality images of *A. thaliana* root systems (Figure 3C).

**Figure 3:**
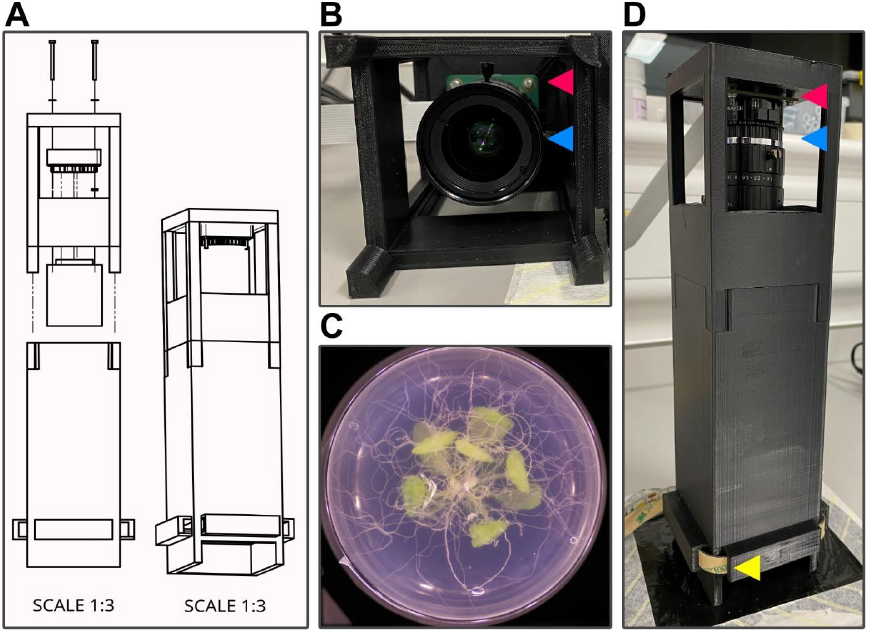
The imaging tower. A) An explosion diagram of the assembly and final product of the imaging tower on scale of 1:3. B) a look inside the top half of the imaging tower. C) An example image taken using the imaging tower. D) The assembled imaging tower. The lens (blue arrow) is mounted to the camera (red arrow), which is connected to the Raspberry Pi (not shown). Consistent all around lighting is provided by an LED strip (yellow arrow).

Printing the imaging tower takes 30 hours (on a Ultimaker S3, Ultimaker PLA with 20% infill), with 30 minutes hands on time for assembly of the device. Imaging efficiency may vary depending on the speed of the Raspberry Pi and if cooling is provided. In our hands, imaging takes approximately 7 seconds per capture. Using these images, the extent and nature of the root system pre-infection can be readily and non-destructively analysed (Figure 4A).

**Figure 4:**
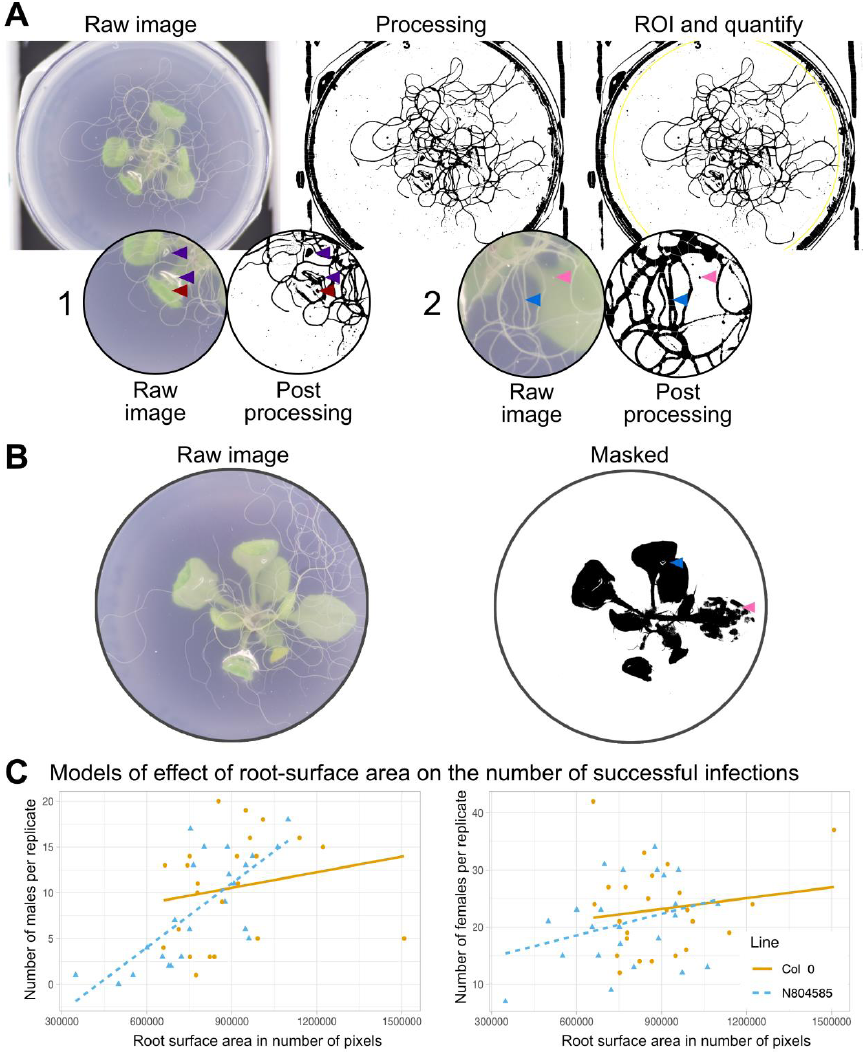
Extraction of root surface area from images and correlation with nematode counts. A) A visual representation of the pipeline used for calculating the root surface area. From left to right the features are extracted from the image and a region of interest is defined. Artefacts picked up in the pipeline: A1) Damage to the agar (purple arrow) and a leaf touching the agar (brown arrow). A2) A small particle on the surface of the plastic (pink arrow), clustered roots interpreted as a singular root (blue arrow). B) an example of the ability to extract the leaf surface area using colour thresholding. Artifacts are introduced by difference in leaf colour (pink arrow) and damage to the agar (blue arrow). C) Two models describing the relationship between number of nematodes and root surface area for two A. thaliana genotypes. The models are described as follows; Columbia 0, Males: y = 5.4+5.7*10^−6^x, Females: y = 17.5+6.33*10^−6^x. N804585, Males: y = 10+2.34*10^−5^x, Females: y = 11+1.26*10^−5^x.

By means of thresholding, a technique that groups pixels into distinct classes (typically a foreground and a background), the root surface area for *A. thaliana* grown in tissue culture was estimated from an image given in number of pixels (Figure 4A). Using this technique most roots can be isolated and quantified. Minor artefacts were introduced (leaves touching the agar, damage to the agar (Figure 4-A1), and dust on the outside of the plates (Figure 4-A2) but do not contribute substantially to the total root surface area and, importantly, are consistently present on all plates, and therefore of minimal concern. A similar method of colour thresholding was used to extract the leaf surface area from an image (Figure 4B), with similar constraints.

### The impact of root surface area on infection

Two genotypes were compared; Columbia 0 and a T-DNA knockout mutant for AT1G07540 (N804585), a plant gene highly expressed during nematode infection [30]. Using these estimates of root area from the images, the relationship between the number of nematodes and the total root surface area for the two genotypes has been summarised by the linear models shown in Figure 4C. The number of manually counted nematodes for both male and female increased with the total root surface area at inoculation density of 80 infective stage nematodes/plant. Approximately 4% of female and 3% of male variation in nematode number on Colombia wild-type and 10% of female and 54% of the variation in males on mutant line N804585 can be explained by root area pre infection (Figure 4C). Together, the positive correlation in Figure 4C demonstrates the importance of normalising nematode counts by available root are pre-infection, when comparing between but even within genotypes.

### Validation of image analyses method of cyst nematode infection

The image quality from the imaging tower not only allows for quantification of roots, but also clearly shows female life stage nematode on roots from 21 days post infection (dpi) onwards (16-hour day at 21°C and 8-hour night at 20°C). There is, therefore, scope to employ similar image analysis scripts for the detection of nematodes directly from images of infected plants.

### Counting cysts

Like detecting roots from images, high contrast between the entity of interest and the background is crucial for threshold-mediated object isolation. Once extracted, a feature can be scored for various parameters. In images, the encysted *H. schachtii* often has excellent contrast with the background, due to its tanned cuticle, and was the initial life stage of interest for automated analyses. For a series of images, each containing a single infected plant, the colour histograms were normalised to an arbitrary reference (any one of the series) using a custom python script [31]. The optimal colour thresholding for the reference image was then empirically determined to isolate the cysts, and then applied to all other images in the series. Using this technique, 147/230 cysts were identified, with 3 false positives (1.3 %) introduced by artefacts on the petri-dish. Of those cysts that were identified, their area was quantified automatically. Taken together, cysts were surprisingly difficult to identify and quantify using threshold-mediated object isolation.

### Counting females

In order to identify females of *H. schachtii* (i.e. before encystment) using threshold-mediated object isolation, it was anticipated that some additional steps would be required to increase contrast: at this life stage, they are very similar in colour to roots and senesced *A. thaliana* leaves. In standard KNOP media, we were unable to identify parameters that would distinguish females from surrounding roots (data not shown). To address this, we tested various contrasts by dyeing the agar with food colouring, followed by illumination with white, red, green or blue lights (Figure 5A, B). The best contrast was found using a red food colouring (Limino, DYL-ghm-20210126-324) illuminated with blue light (380-500 nm, Figure 5B). Green light provides a similarly good contrast but was more prone to highlighting artifacts (data not shown).

**Figure 5:**
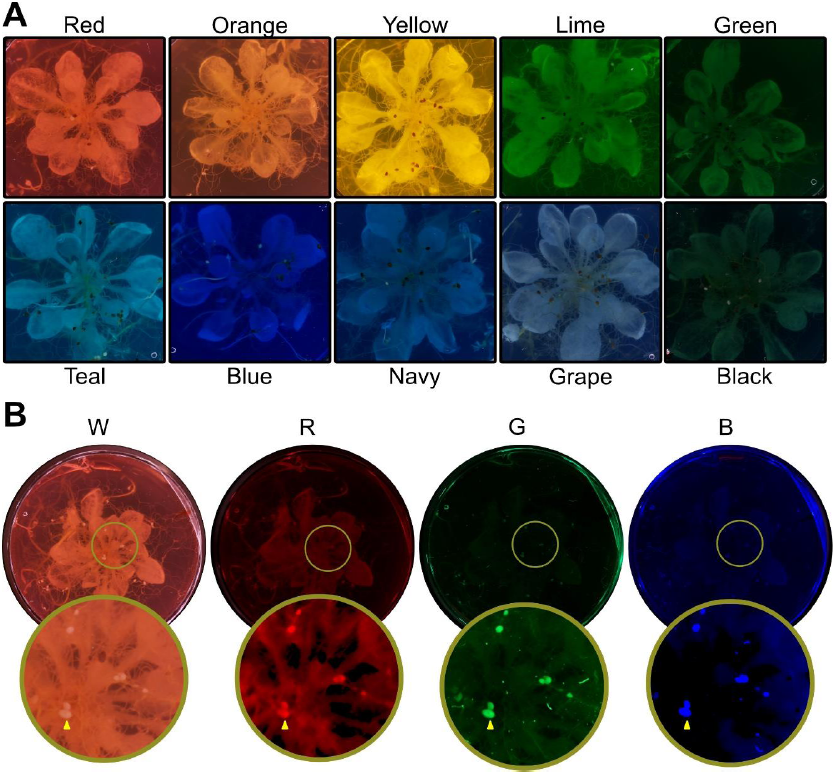
The spectrum of coloured agar under various lighting conditions. A) The range of colours tested for contrast with nematodes and filtration potential of the background. B) The colour red agar under four different lighting conditions. From left to right: White, Red, Green and Blue light. The blue lighting condition with the red agar were the most ideal combination for quantifying nematodes using the segmentation tool. The bottom image row has the contrast and brightness adjusted for visibility. Examples of female nematodes are highlighted with yellow arrows.

Using a well-established ImageJ library (morphological segmentation [29]) and manual verification, we extracted nematodes from images with 93.3% (70/75) accuracy. We were also able to measure nematode area from the images, with an R^2^ of 0.83 with manual area quantification compared to only an R^2^ of 0.44 when counting cysts (Figure 6D). The automated segmentation tool was not able to reliably extract cysts from these conditions. For validation of these methods, the outcomes of automated and manual area were compared against each other (Figure 6D).

**Figure 6:**
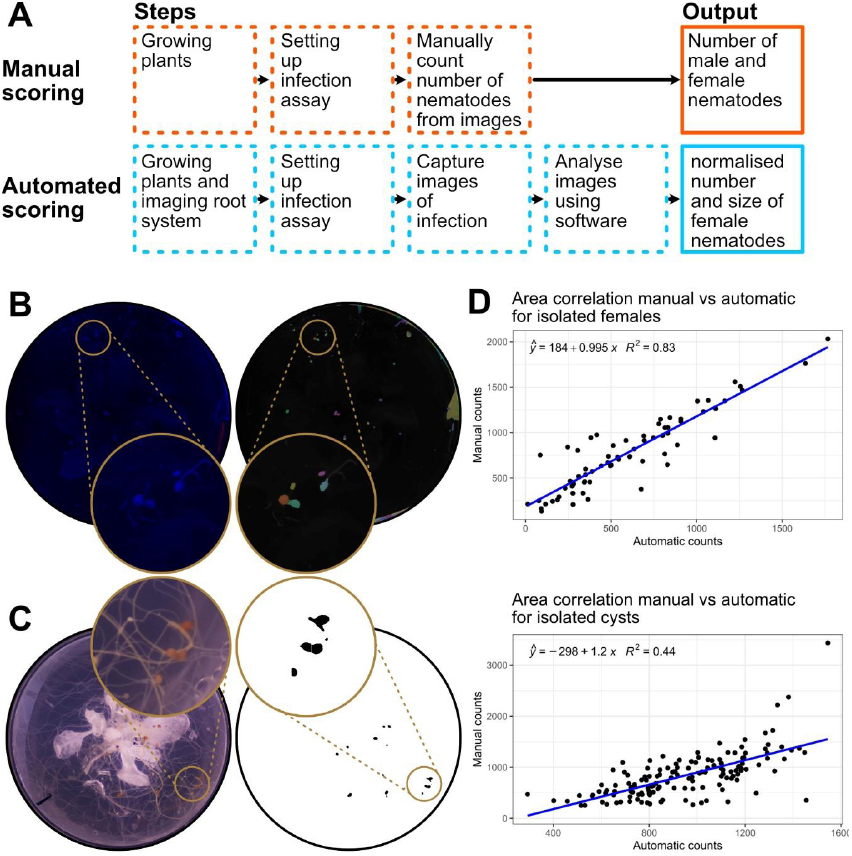
Isolation and quantification of nematodes from images. A) A schematic workflow highlighting the steps required for manual and automated nematode counting, and the differences in output. B) Isolation of female cyst nematodes from red coloured agar under blue light illumination, and the extracted overlay from the script. C) Isolation of encysted nematodes under white light illumination, and the extracted overlay from the script. D) Correlation of area of the nematode between manual counts and automatic counts using the script for females and cysts.

## Discussion

Our knowledge on both resistance genes and susceptibility genes against plant parasitic nematodes remains scarce. The most widely studied, Gpa2, recognises the nematode effector RBP-1 once it is delivered into the syncytial feeding site: triggering an immune response and ultimately resistance [32]. For *Heterodera schachtii*, a commonly used model species for cyst nematodes, resistance associated with HS1^pro1^ was introduced in sugar beet through introgressive hybridization [33]. The presence of this gene diminishes the growth of the feeding organ, impacting the reproduction of the nematode [34]. Despite the positive effects on nematode population control in the field, the introgressed gene reduces yield [35], and is therefore not widely used [36]. However, control is not limited to resistance, the absence of susceptibility genes can also determine the outcome of infection illustrated by the deletion of HIPP27 and the more well understood vitamin B5 pathway gene AtPANB1 [13,37]. A broad focus of the field is to identify genes that have direct impact on nematode parasitism.

The *Heterodera schachtii:Arabidopsis thaliana* infection assay is broadly used for screening of genes due to the availability of the genetic resources on the plant side of the interaction. One of the major bottlenecks that remains in the phenotyping assay, it: i) requires access to costly microscopy equipment; ii) lacks the ability to readily account for phenotypic variation of the infectable tissue of the host; and iii) requires considerable time investment of a trained operator. Taken together, only a relatively small number of genotypes can be screened.

To address the first constraint, a cheap, easy to build and use, 3D-printed imaging tower was created. The construction consists of few parts ensuring ease of assembly and, with the growing number of 3D printing services [38], is available at low cost (printing service ∼£67.56 with delivery [39], or £10.87 in house at time of writing). The assembly can take images of petri-dishes every seven seconds. At two weeks post germination, the root system of *A. thaliana* can be isolated from the photographs using a threshold-based image analyses method and quantified in a measure of pixels.

It is hypothesised that not only the genotypic, but also the phenotypic variation of the host impacts its infectability. For example, some loss of function mutants vary in their susceptibility or resistance to nematodes [40], but in some cases also vary in their root physiology, complicating the assessment. It is hypothesised that the available root area before infection acts as a direct physical constraint to cyst nematode parasitism, which can affect the outcome of a susceptibility assay.

Due to the high variability in nematode infection a good model remains difficult to construct. The linear model in Figure 4, however, illustrates an interaction between the root surface area and the number of nematodes. Overall, the number of nematodes increases with the amount of root surface area for a given inoculum. Importantly, between the two tested genotypes there is a clear difference in magnitude of the effect.

On images of the same dishes at a later stage (21+ days post infection), both female and cyst life stage are clearly visible. A fully automated script for counting cysts using colour thresholding from petri-dishes was created. This method, while quick, has a high error rate (63.9 % accuracy, 1.3 % false positive and 34.8 % false negative). Adjusting the threshold can reduce the number of false negatives, however, this typically increased the number of false positives. The technique is largely limited by the complexity of the background around nematodes and the variability in colour and contrasts between plates. To mitigate against this, various colour dyes were added to the agar which, under blue illuminated light, increase the contrast of female nematodes, and weaken the visibility of the background. The greatest distinction between nematode and background was found using Limino red food colouring illuminated with blue light (380-500 nm). Using both a program which isolates features from an image by means of morphological operations, and manual verification, the frequency and size of the nematode could be determined with a 93.3 % accuracy and 6.7 % false negatives. In addition to count, this method provides a good size correlation between fully manual and semi-automatic measurements (R^2^ = 0.83). Importantly, from the available data, it is unclear whether the dye is entirely innocuous (some dyes did inhibit germination, or root growth, and were not included in the experiment); regardless the impact of dye can be disregarded with consistent use between the control and the treatment. Some coloured dyes do contribute to the false positives by highlighting particles on the outside of the petri-dish which must be filtered out manually. Through modifications to the traditional assay, a semi-automatic and high accuracy measurement of infection can be achieved. This, however, does little to reduce the effort involved for screening females as manual verification of the isolated mask is required. Regardless, at no extra effort, the biologically relevant measurement of nematode size is gained, which makes the semi-automated screening superior over manual screening for females.

A major limitation of this new screening method is that it does not allow for inclusion of male nematodes. Whilst visible on images of non-coloured agar, the current pixel density does not create a sharp enough image to reliably distinguish them from the background. This limitation may be addressed in the future with a higher resolution camera and a more appropriate lens allowing for reduction of undesired imaging of the area around the petri dish, optimising the use of available pixels. Furthermore, the new technique is limited by its throughput as it does not allow for a decrease in time required for screening. An issue that may be addressed in the future by further automation on the hardware side.

These first steps towards digitalizing screening are crucial in the development of a fully automated system for screening of plant parasitic nematode infection. The main limitation of the protocol to date is speed, and the requirement of modification of the agar using coloured dyes for accuracy of automated screening. Classical computer vision techniques are restricted in their capabilities to isolate objects from complex backgrounds. These traditional methods, however, can be accompanied by deep learning approaches to lift these constrains [41], and have proven already valuable in phenotyping of plant diseases [42]. Ideally, this would lift the requirement of modified agar, and manual verification of nematode count, essentially limiting the speed of screening to the speed of imaging.

## Conclusion

Overall, this paper describes an accessible open-source image-based method for better insight in cyst nematode infection on *Arabidopsis thaliana* by including the size of the nematode as a parameter in infection screening, normalised for available infectable tissue. Further research is needed to elucidate the exact interaction between root surface area and the success on nematode infection. While the new method adds a new dimension to nematode infection screening, it is still limited in speed. Beyond the scope of this paper, machine learning approaches may lift the constraints of manual verification in the assay, and thus further increase screening speed.

## Author Contributions

O.P.K., designed the hardware and software.

O.P.K., I.K., R.H., U.S., S.W., B.S., and F.D.B. collected the data.

O.P.K., J.Z., and S.E.v.d.A wrote the main manuscript.

O.P.K. prepared the figures.

All authors reviewed the manuscript.

## Acknowledgments

Work on plant-parasitic nematodes at the University of Cambridge is supported by DEFRA licence 125034/359149/3, and funded by BBSRC grants BB/R011311/1, BB/N021908/1, and BB/S006397/1. This work is also supported by an overseas foreign specialist project from the Ministry of Science and Technology of the People’s Republic of China: G2021145005L. For the purpose of open access, the author has applied a Creative Commons Attribution (CC BY) licence to any Author Accepted Manuscript version arising from this submission.

